# Evolutionary and geometric signatures reveal ligand-binding sites across proteomes

**DOI:** 10.1101/2025.10.07.680847

**Authors:** Stelina Tarasi, Laura Malo, Alexis Molina

**Affiliations:** Department of Artificial Intelligence, Nostrum Biodiscovery, Barcelona, 08029, Spain; Department of Drug Discovery, Nostrum Biodiscovery, Barcelona, 08029, Spain

**Keywords:** binding site prediction, cryptic sites, protein language model, protein structure representation

## Abstract

Identifying protein binding sites is central to drug discovery, yet many computational approaches still trade off precision, recall, or throughput when scaled. We introduce PickPocket, a deep learning model that fuses evolutionary information from protein sequences with geometric representations of structure to identify ligand-binding residues at proteome scale. By leveraging complementary sequence context and spatial neighborhoods, PickPocket generalizes across diverse protein families and ligand chemistries while operating at a recall-oriented setting with competitive precision. In benchmark evaluations it delivers strong residue-level recovery and, despite no explicit training on conformational switching, reliably identifies cryptic pockets in held-out structures, comparing favorably with specialized approaches. Applied across 356,711 proteins, the method nominates previously unannotated candidate sites enriched for functional signals and highlights tractable surface chemistry on therapeutically relevant targets. These results position evolutionary-geometric fusion as a practical foundation for large-scale site mapping that can shorten the path from structure to experiment and support hit discovery, mutagenesis design, and target assessment.

## 1 Introduction

Protein function is often modulated by small molecules or peptides that bind to specific surface regions, altering structural dynamics and biochemical activity [1]. Accurately identifying these binding sites is a central step in rational drug discovery since it guides molecular design and supports target selectivity [2]. Classical computational strategies, i.e. molecular docking, geometric pocket detection, and physicochemical mapping, remain useful, but they frequently need prior knowledge of the binding region and struggle with cryptic or allosteric sites that emerge only under specific conformational states [3, 4].

Geometric and biophysical methods identify and characterize pockets from structure alone. For example, fpocket uses Voronoi tessellation and alpha spheres to delineate cavities which are then ranked using geometric and physicochemical descriptors [5]. LIGSITE, building on POCKET, scans cubic grids to score Protein–Solvent–Protein (PSP) events along axes and diagonals, reducing orientation sensitivity while remaining fast enough for large-scale screening [6, 7]. PocketFinder contours smoothed Lennard–Jones potential maps to produce ligand-binding predictions that are comparatively robust to modest conformational changes [8]. Despite their efficiency and interpretability, these methods can be limited by their reliance on a single (often holo) structure and by the difficulty of capturing long-range cooperativity.

Learning-based approaches augment or replace hand-crafted rules with data-driven features. Early machine-learning pipelines such as PRANK and P2Rank re-rank or predict ligandability from local surface chemistry and geometry, delivering strong speed–accuracy trade-offs [9, 10]. Deep learning models extend this trend as 3D CNNs like PUResNet voxelize proteins and use residual U-Net architectures with Dice-style objectives to address class imbalance on curated sc-PDB splits [11–13]. Graph neural networks (GNNs) bring rotational equivariance and improved message passing; VN-EGNN introduces virtual nodes to better represent pocket geometry and mitigate oversquashing, while GrASP leverages attention to fuse atomic and residue contexts for precise ligandable-atom predictions [14–18]. More recently, protein language models (pLMs) have enabled structure-aware yet evolution-informed predictors. IF-SitePred couples ESM-IF1 embeddings with LightGBM and DBSCAN clustering to localize sites from backbone geometry, showing resilience on AlphaFold2 inputs [19–22].

However, important gaps remain. Generalising from bound to unbound states, coupling evolutionary conservation with 3D geometry, and scaling reliably to proteome-wide inference. To address these limitations, we introduce PickPocket, a binding-site predictor that integrates evolutionary and geometric signals by combining ESM-2 embeddings [23] with GearNet-based graph representations of protein structure [24]. This fusion captures sequence-derived functional constraints and spatial residue–residue relationships, enabling robust identification of conserved, functionally relevant cavities, including cases where side-chain placement or conformational state is uncertain. We show that PickPocket outperforms existing methods on relevant benchmarks and scales to proteome-wide deployment, offering a practical tool for target assessment and early ligand discovery.

## 2 Results and Discussion

### 2.1 PickPocket Effectively Learns to Predict Binding Pockets

PickPocket was evaluated against state-of-the-art binding site prediction methods using the LIGYSIS [25] benchmark, which provides a comprehensive dataset of ligand-binding sites across diverse protein structures. Unlike previous benchmarks, LIGYSIS aggregates binding interfaces across multiple protein structures, includes diverse ligands, and prioritizes biological units for functional relevance. After filtering chains with missing UniProt [26] mappings, the final set comprises 2,775 protein chains. To meet ESM-2 embedding constraints, proteins over 1,022 residues were cropped, excluding 7 pockets from evaluation.

For residue-level classification, we used standard metrics, specifically the F1 score and Matthews Correlation Coefficient (MCC), to evaluate whether residues belong to a binding site. The results indicate that PickPocket achieves the highest F1 score (0.42) and maintains a competitive MCC of 0.37, outperforming existing deep learning-based methods such as PUResNet, GrASP, and P2Rank CONS [27], as well as classical geometric approaches like fpocket, PocketFinder, and Ligsite (Table 1). These findings highlight PickPocket’s ability to balance precision and recall, making it a robust approach for identifying functionally relevant binding pockets.

**Table 1:** Performance comparison ordered by F1 score.

| Method | F1 | MCC |
| --- | --- | --- |
| PickPocket | **0.42** | 0.37 |
| PUResNet | 0.41 | **0.39** |
| GrASP | 0.39 | 0.34 |
| P2Rank_CONS | 0.36 | 0.30 |
| P2Rank | 0.31 | 0.26 |
| PocketFinder | 0.31 | 0.22 |
| Ligsite | 0.31 | 0.21 |
| VN-EGNN | 0.29 | 0.26 |
| IF-SitePred | 0.29 | 0.24 |
| Surfnet | 0.29 | 0.20 |
| DeepPocket_SEG | 0.27 | 0.21 |
| fpocket | 0.23 | 0.12 |

To assess its effectiveness in binding site detection, PickPocket was evaluated using Top-N recall as a metric. Top-N recall measures the model’s ability to identify true binding sites among its top-ranked predictions. A prediction is considered correct if the distance between the predicted pocket centroid and the observed binding site centroid (DCC) is ≤ 12 Å. When evaluating recall, we consider both the Top-N and Top-(N+2) ranked predictions, where N is the number of true binding sites in the protein. Following the approach used in LIGYSIS, we compute the recall as:

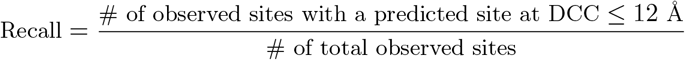

PickPocket achieves a Top-N recall of 48.9%, the highest among the benchmarked methods, outperforming state-of-the-art approaches such as P2Rank, GrASP, and DeepPocket. Additionally, it maintains a Top-N+2 recall of 53.3%, demonstrating consistent performance across different ranking cutoffs (Table 2). These results highlight PickPocket’s superior ability to retrieve relevant binding sites when benchmarked against established methods.

**Table 2:** Recall performance ordered by Top-N recall.

| Method | Top-N | Top-N+2 |
| --- | --- | --- |
| PickPocket | **48.9** | 53.3 |
| P2Rank_CONS | 48.8 | 53.9 |
| fpocket_PRANK | 48.8 | **60.4** |
| GrASP | 48.0 | 49.9 |
| P2Rank | 46.7 | 51.9 |
| DeepPocket_RESC | 46.6 | 58.1 |
| Ligsite_AA | 44.9 | 49.0 |
| VN-EGNN_NR | 44.5 | 46.1 |
| PocketFinder_AA | 44.5 | 48.9 |
| DeepPocket_SEG-NR | 43.4 | 49.4 |
| Surfnet_AA | 43.3 | 47.4 |
| PUResNet_PRANK | 40.8 | 41.1 |
| fpocket | 38.8 | 46.5 |
| IF-SitePred_RESC-NR | 29.7 | 39.1 |

**Table 3:** Overall ranking of methods grouped by core method names (best rank per group), ordered by total rank.

| Method Group | Top-N | Top-N+2 | F1 | MCC | Total |
| --- | --- | --- | --- | --- | --- |
| PickPocket | **1** | 4 | **1** | 2 | **8** |
| P2Rank (incl. CONS) | 2 | 3 | 4 | 4 | 13 |
| GrASP | 4 | 5 | 3 | 3 | 15 |
| PUResNet (incl. PRANK) | 10 | 10 | 2 | **1** | 23 |
| fpocket (incl. PRANK) | 2 | **1** | 11 | 11 | 25 |
| DeepPocket (incl. SEG, RESC) | 5 | 2 | 10 | 8 | 25 |
| Ligsite (incl. AA) | 6 | 6 | 5 | 8 | 25 |
| PocketFinder (incl. AA) | 7 | 7 | 5 | 7 | 26 |
| VN-EGNN (incl. NR) | 7 | 9 | 7 | 5 | 28 |
| Surfnet (incl. AA) | 9 | 8 | 7 | 10 | 34 |
| IF-SitePred (incl. RESC-NR) | 11 | 11 | 7 | 6 | 35 |

**Table 4:**
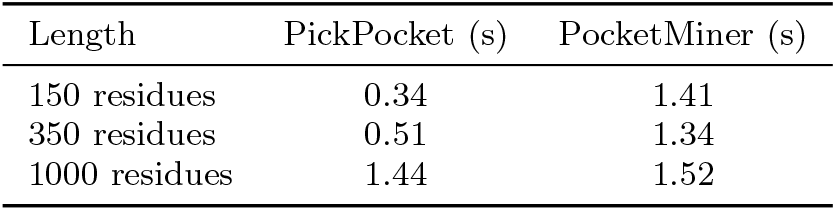
Total CPU Inference Time Comparison.

| Length | PickPocket (s) | PocketMiner (s) |
| --- | --- | --- |
| 150 residues | 0.34 | 1.41 |
| 350 residues | 0.51 | 1.34 |
| 1000 residues | 1.44 | 1.52 |

**Table 5:** Summary of PickPocket recall on experimentally defined pockets. Recall is the fraction of experimentally validated residues recovered by the model.

| Protein system | Pocket / Site | Pred. / Total | Recall (%) |
| --- | --- | --- | --- |
| DAT | S1 orthosteric | 5 / 6 | 83 |
| DAT | S2 vestibular | 1 / 4 | 25 |
| SV2A | TeNT-HC interface | 0 / 5 | 0 |
| SV2A | LEV pocket | 13 / 13 | 100 |
| hRpn13 | Pru-A core | 3 / 4 | 75 |
| hRpn13 | Pru-B | 1 / 4 | 25 |
| CB1 | VIP36 cryptic site | 2 / 5 | 40 |
| FFA2 | Orthosteric | 10 / 13 | 77 |
| FFA2 | Allosteric 1 | 7 / 13 | 54 |
| FFA2 | Allosteric 2 | 0 / 6 | 0 |
| IL-36R | D1 orthosteric | 10 / 12 | 83 |

Compared to other methods, PickPocket benefits from a combination of protein language models and graph-based structural learning. The integration of ESM-2 embeddings allows it to incorporate evolutionary information, which is known to be relevant for binding site detection. At the same time, the use of GearNet enables it to capture spatial relationships between residues, complementing the sequence-based information. This dual approach helps improve both recall and precision by identifying functionally relevant binding sites while reducing the likelihood of detecting non-functional surface cavities.

### 2.2 PickPocket Predicts Cryptic Sites Without Specific Training

Identifying binding pockets that are not immediately apparent from static protein structures remains a significant challenge in drug discovery. Many functionally relevant sites, such as cryptic pockets that emerge upon ligand binding or allosteric sites that regulate protein activity from a distance, are often missed by traditional structure-based methods. These pockets may be transient, hidden in unbound structures, or only form under specific conformational states. Accurately detecting such sites requires a model capable of integrating sequence-derived evolutionary information, structural flexibility, and binding-relevant physicochemical features.

To evaluate PickPocket’s ability to predict cryptic binding sites, we analyzed 24 apo structures from the PocketMiner test set and compared its predictions to the corresponding holo structures. Unlike PocketMiner, which is explicitly trained to detect cryptic pockets using molecular dynamics derived labels, PickPocket operates without direct supervision for cryptic site prediction.

Quantitative assessment of PickPocket’s capability in identifying cryptic pockets was performed using Precision-Recall (PR) curves, comparing its performance against PocketMiner (Figures 3a and 3b). PickPocket consistently outperformed PocketMiner in both apo and holo structures, achieving significantly higher AUC scores (0.617 and 0.659) compared to PocketMiner’s (0.438 and 0.529), respectively.

**Fig. 1:**
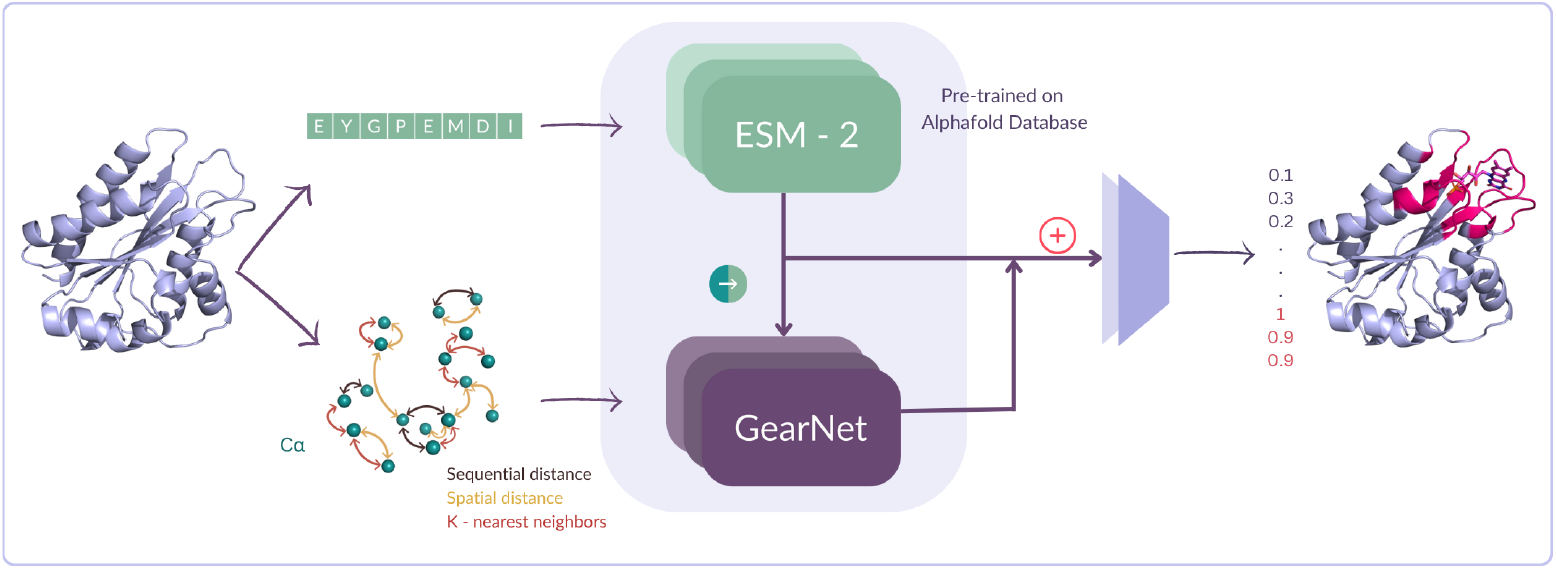
PickPocket architecture for protein binding site prediction. The model combines sequence embeddings from ESM-2 with structural graph features processed by GearNet through serial fusion. ESM-2’s contextual representations initialize GearNet’s node features, integrating evolutionarily conserved sequence patterns with geometric structural information. The fused embeddings are passed to a classification head that outputs the probability of each residue belonging to a binding site.

**Fig. 2:**
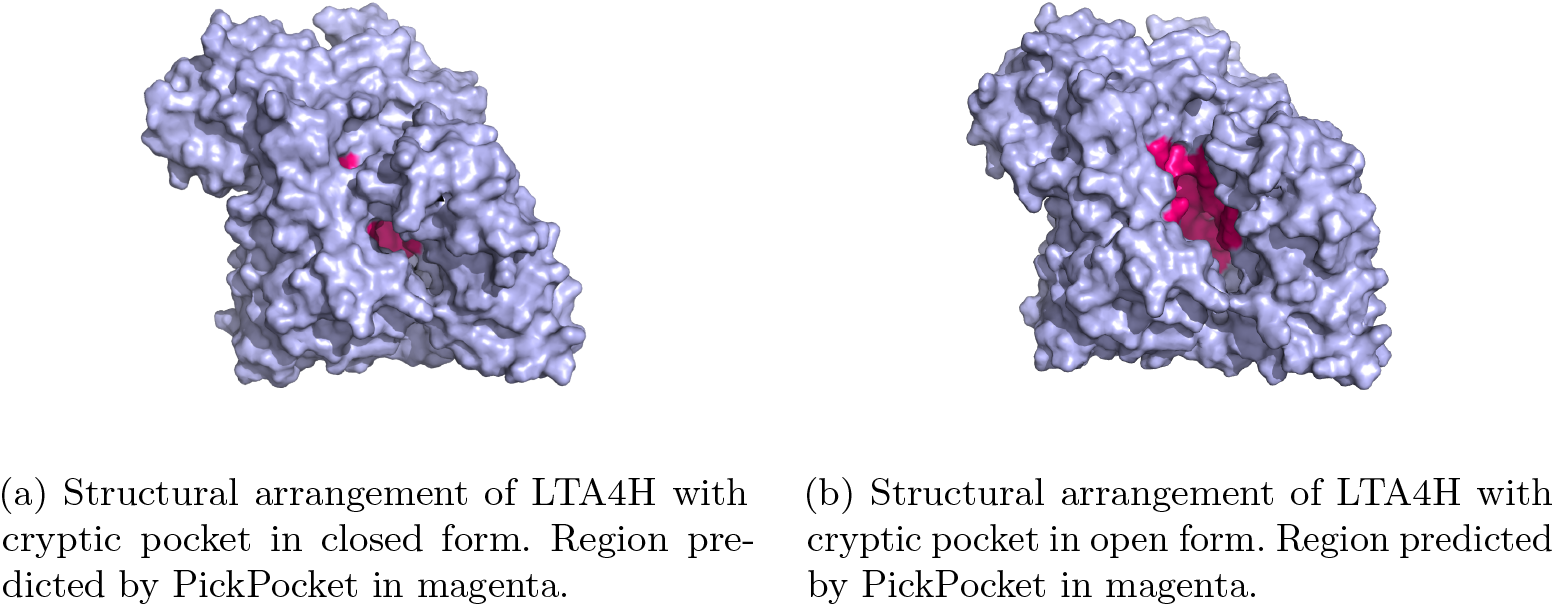
PickPocket accurately identifies binding sites in both conformations: (a) closed pocket (PDB: 5NI6) and (b) open pocket (PDB: 5NIA).

**Fig. 3:**
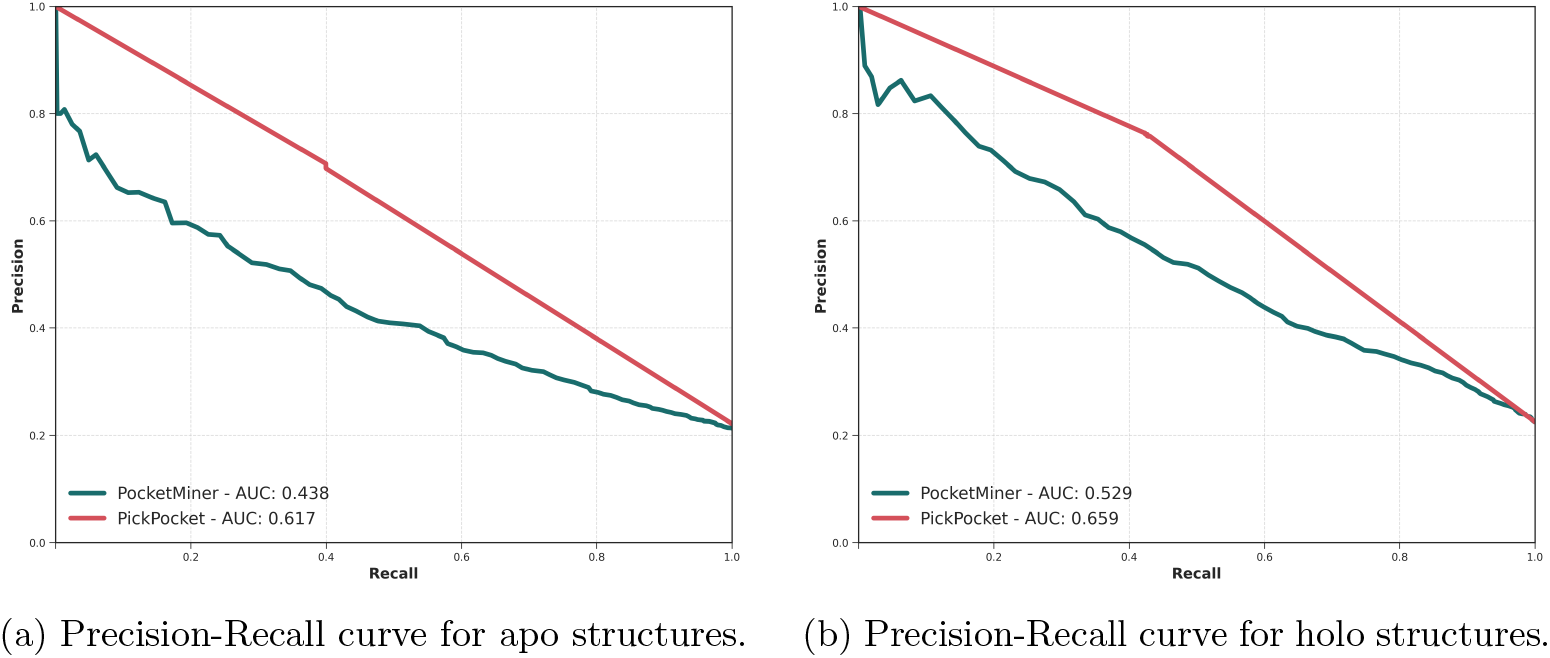
Comparison of PickPocket and PocketMiner on apo (unbound) and holo (bound) structures for cryptic pocket prediction. PickPocket consistently achieves superior performance, generalizing across different protein conformations without requiring MD-derived labels.

The PR curves highlight that PickPocket maintains superior precision across a broader recall range, particularly in the apo structures (Figure 3a), where cryptic pockets are inherently more challenging to detect due to the absence of ligand-induced conformational rearrangements. The improved AUC suggests that PickPocket effectively captures latent structural and evolutionary signals that correlate with cryptic site formation, even in unbound states.

In the holo structures (Figure 3b), where ligand-induced conformational changes reveal binding pockets more explicitly, PickPocket continues to outperform PocketMiner. Its higher AUC (0.659) indicates that it is capable of accurately localizing binding pockets even in cases where the pocket has undergone a cryptic-to-open transition. The consistency in its performance across both apo and holo structures suggests that PickPocket generalizes well to cryptic pocket identification without requiring explicit cryptic pocket training.

This underscores the scalability and efficiency of PickPocket for proteome-wide cryptic site discovery, offering a valuable alternative to MD-based approaches while significantly reducing computational costs.

Moreover, we performed an embedding-level analysis using centered kernel alignment (CKA) to quantify apo–holo shifts in residue representations. Across four apo–holo pairs, pocket residues showed lower similarity between states than non-pocket residues (mean ΔCKA = − CKAsite CKAnon = − 0.133 ± 0.103; three of four pairs negative), indicating that PickPocket’s embeddings change more where cryptic rearrangements occur while remaining stable elsewhere. This supports sensitivity to functionally relevant conformational change rather than overfitting to static geometry. Implementation details, residue matching, alignment criteria, and robustness checks are in Appendix B.

### 2.3 Validation on Newly Resolved Experimental Structures and Binding Sites

To assess out-of-distribution generalisation and practical utility, we assembled a panel of recently reported protein–ligand complexes whose binding sites were mapped experimentally by X-ray, cryo-EM, or NMR, then (i) downloaded the deposited PDB coordinates, reconciled residue numbering with the canonical sequence used in the primary publication (accounting for signal-peptide truncations, affinity tags, or ectodomain deletions), and (iii) compiled the residues supported by mutagenesis or density-based contact analysis as directly contributing to ligand binding, PickPocket was run on raw atomic coordinates with no ligand or prior site information, residues were called positive at binding probability *>* 0.7, positives were clustered into pockets with DBSCAN, and for each experimentally defined site we computed residue-level recall (predicted / total true) while annotating systematic misses such as surface-exposed, highly flexible, or transiently interacting side chains.

Across seven newly resolved systems, the outward-open human dopamine transporter DAT (PDB 9EO4) [28], synaptic vesicle glycoprotein 2A SV2A in levetiracetam-bound and TeNT-HC interface conformations (PDB 9FYR) [29], the proteasome receptor subunit 13 hRpn13 bound to ENT (PDB 8VWO) [30], the TBXT transcription factor in apo/holo states (PDBs 6F58/8CDN) [31], the active VIP36-bound CB1 receptor (PDB 9B54) [32], free fatty-acid receptor 2 FFA2 in active/inactive conformations (PDB 8Y6Y) [33], and the interleukin-36 receptor IL-36R with peptide and small-molecule ligands (PDB 9ETI) [34], a consistent predictive pattern was detected. Through PickPocket, rigid, concave, hydrophobic/aromatic orthosteric cavities are recovered with high fidelity, whereas shallow, flexible, electrostatic, or strongly state-dependent allosteric sites are more challenging.

Concretely, DAT’s orthosteric S1 pocket reached 83% recall with D79, V152, F320, S321, and S422 recovered and Y88 missed, likely due to a flexible boundary-layer position, while the vestibular S2 pocket reached 25% (G386 captured; solvent-exposed K92, K384, H477 missed), though a contiguous high-score cluster remained aligned with the verified region, indicating partial recognition of the shallow interface (Figure 4a). In SV2A, the flat *β*-strand augmentation mediating TeNT-HC binding was essentially missed, consistent with a concavity preference, whereas the deep, aromatic-rich levetiracetam pocket was perfectly recovered (100%) with neighbouring high-score residues suggesting a plausible cavity extension. For hRpn13–ENT, the primary Pru-A pocket achieved 75% recall (P40, S90, W108 recovered; solvent-exposed K42 missed), while the shallow, electrostatic Pru-B site reached 25%, with additional hydrophobic clusters hinting at broader interaction pathways. In CB1’s cryptic VIP36 site the model captured geometrically/electrostatically prominent determinants, D163^2.50^ as the ionic hub and W356^6.48^ as a gate, but missed transient or flexible contributors such as N389^7.45^ and F200^3.36^, underscoring the value of modelling conformational plasticity (Figure 4b). In FFA2, performance was strong in the orthosteric pocket (77%) and moderate in allosteric site 1 (54%), but null in allosteric site 2 (0%), attributable to the inactive-state structure lacking the activation-induced cavity and illustrating the need for ensemble or active-state inputs when assessing dynamic sites. Finally, in the IL-36R ectodomain the D1 orthosteric site was recovered at 83% (10/12), including induced-fit/cryptic components, with additional clusters offering plausible routes for ligand optimisation.

**Fig. 4:**
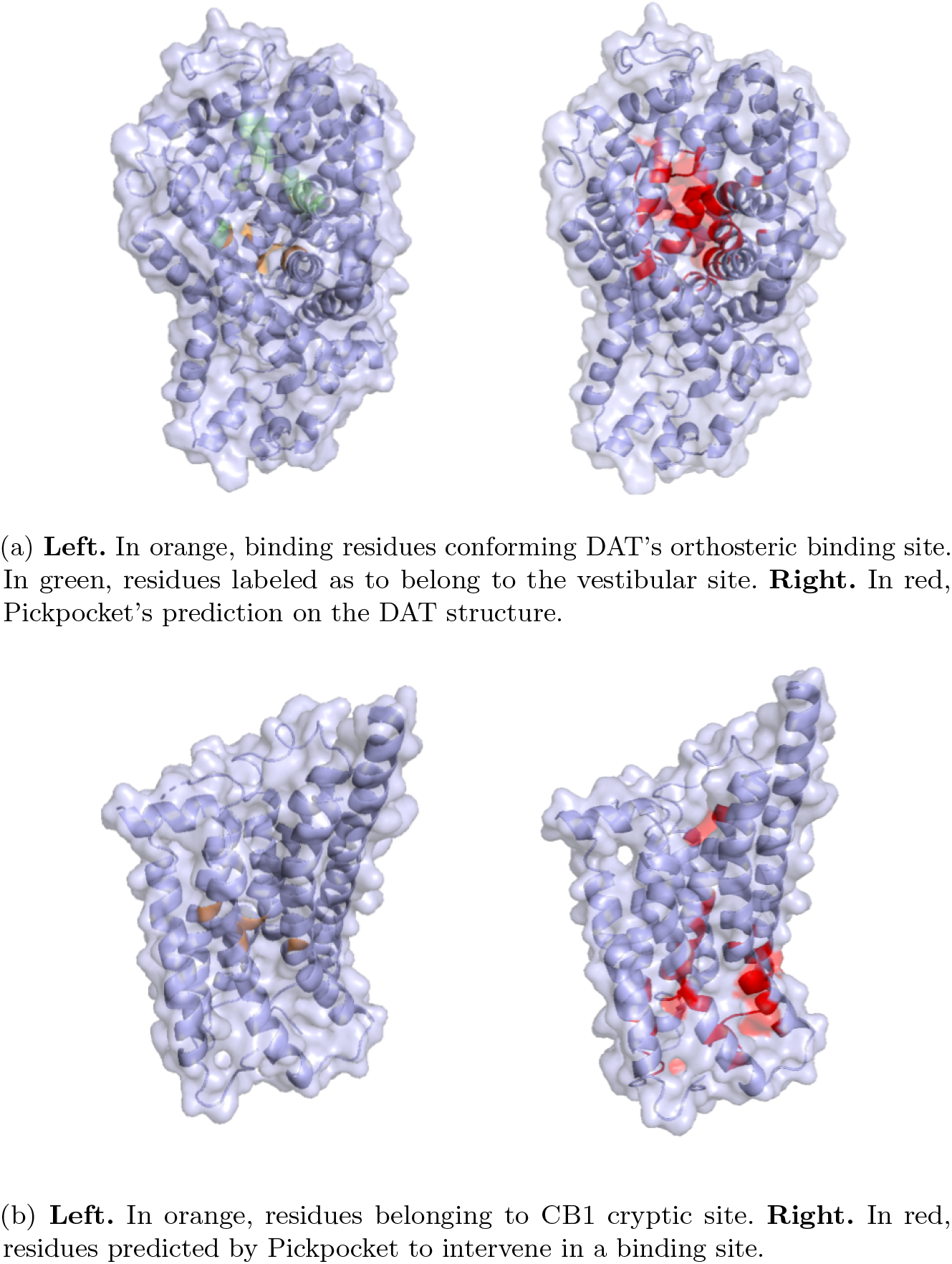
Comparison of predictions and ground truth for DAT (4a) and CB1 (4b).

### 2.4 Evolutionary Information Drives Binding Site Recovery

Here we probe how sequence and structure combine to deliver practical pocket calls and why the recall profile of PickPocket is the right default for discovery. On LIGYSIS, PickPocket attains an F1 of 0.42 with MCC of 0.37 using a residue probability threshold of one half and DBSCAN with epsilon set to 5 angstrom and a minimum of 3 residues. It also recovers the most sites by centroid criteria with Top N recall of 48.9% and Top N+2 of 53.3%, ranking first across F1 and Top N families while remaining competitive on MCC. These results indicate that joint sequence and structure learning retrieves more of the residues that actually line functional cavities than competing approaches that rely on geometry or hand crafted features alone.

Ablations clarify the origin of these gains (Table 6). In our internal variants that keep the ESM and GearNet streams separate at initialization, the Serial and Parallel fusions^1^ produce F1 around 0.38 with MCC around 0.35, trading a little sensitivity for slightly higher global calibration. Single-modality baselines indicate that evolutionary signal carries more weight on this benchmark: ESM alone achieves an F1 of approximately 0.36 and an MCC of 0.30, while GearNet alone performs lower, with an F1 around 0.28 and MCC around 0.25. In the freezing study (Appendix A), freezing ESM causes the largest performance drop, whereas freezing GearNet is less detrimental, indicating a primarily sequence-driven model with geometric refinement. In biological terms, conservation and coevolution guide the model toward patches that tolerate ligand binding across families, while the graph encoder helps consolidate contiguous shells that define cavity extent and chemistry.

**Table 6:**
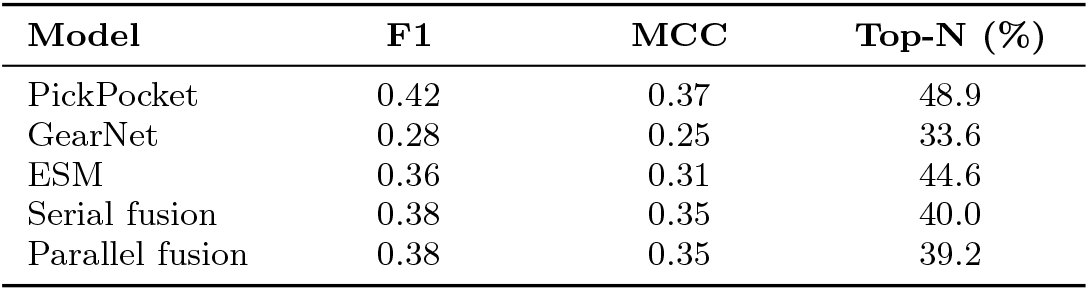
Comparison of model performance metrics for binding site prediction on LIGYSIS dataset. Metrics include F1 score, Matthews correlation coefficient (MCC), and Top-N accuracy (%).

| Model | F1 | MCC | Top-N (%) |
| --- | --- | --- | --- |
| PickPocket | 0.42 | 0.37 | 48.9 |
| GearNet | 0.28 | 0.25 | 33.6 |
| ESM | 0.36 | 0.31 | 44.6 |
| Serial fusion | 0.38 | 0.35 | 40.0 |
| Parallel fusion | 0.38 | 0.35 | 39.2 |

Testing on the PocketMiner cryptic site dataset supports this interpretation. In the apo state (Fig. 5a), ESM, using sequence alone, achieves the highest PR-AUC at 0.644, with PickPocket close behind at 0.617 and GearNet lowest at 0.574, consistent with partially formed pockets where single-snapshot geometry can be anti-signal while evolutionary constraints still localize binding-competent surfaces. In the holo state (Fig. 5b), once the pocket is concealed, PickPocket regains performance (PR-AUC 0.659 vs. 0.652 for ESM and 0.638 for GearNet), indicating that adding spatial context improves over sequence alone. Across both panels, sequence provides the dominant signal and structure refines it, enabling transfer across conformational states without MD-derived supervision.

**Fig. 5:**
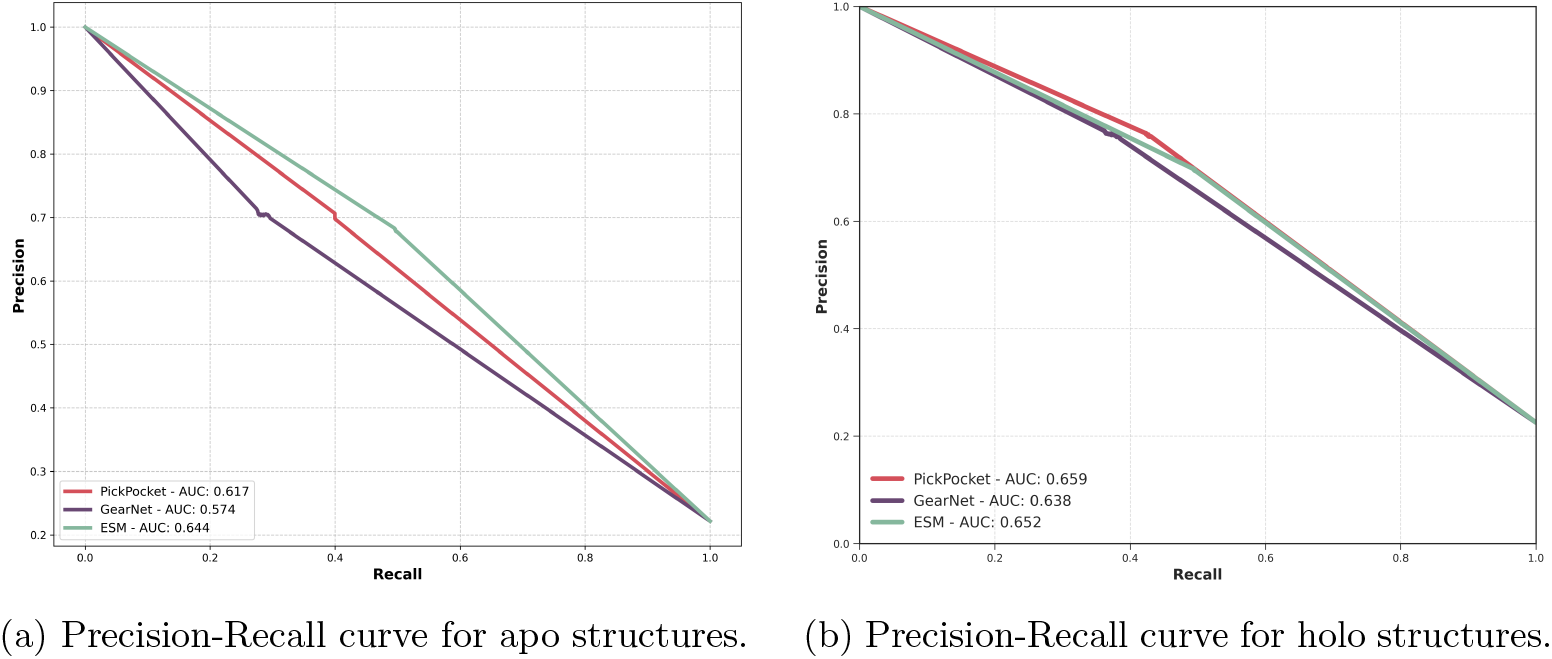
Comparison of PickPocket, ESM and GearNet on apo (unbound) and holo (bound) structures for cryptic pocket prediction.

**Fig. 6:**
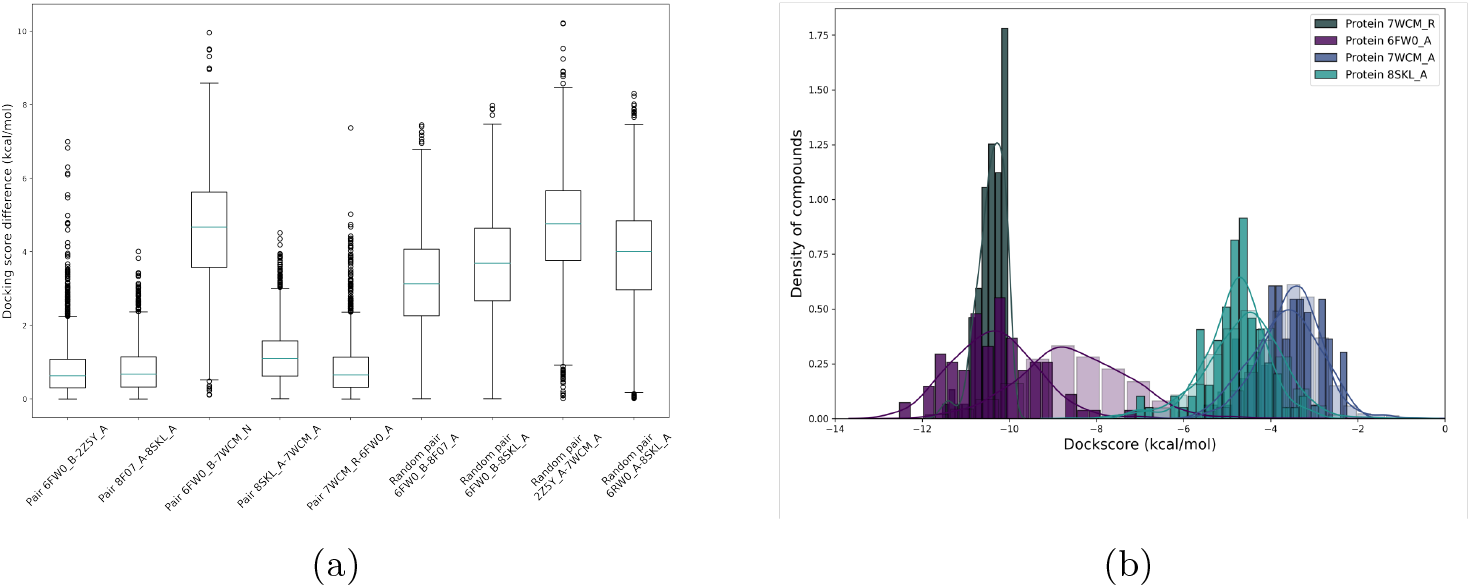
Docking scores for the selected proteins. (a) The docking difference for selected and random pairs, demonstrating that smaller Euclidean distances correlate with more similar docking scores. (b) The distribution of docking scores for the selected pairs, showing a concentration of similar docking scores for these cases. Further details in Appendix C.3

These effects are visible in newly solved systems. Rigid concave hydrophobic or aromatic orthosteric clefts are recovered with high fidelity, exemplified by high recall in the transporter S1 site and the levetiracetam site in SV2A, whereas shallow flexible electrostatic or strongly state dependent allosteric sites remain harder and benefit from conformational ensembles. Where recall is high at the residue level, clustering produces stable pocket calls that seed Top N success, even if a small number of extra surface residues are called positive.

### 2.5 Scaling Binding Site Prediction to All Crystal Protein Structures

To evaluate the scalability and applicability of PickPocket, we predicted ligand-binding pockets across all single-chain protein structures available in the Protein Data Bank (PDB) as of November 2024 [35, 36]. In total, we analyzed 356,711 individual chains, treating each as an independent system. PickPocket identified at least one pocket in 76.2% of chains, with an average of 3.08 pockets per target when at least one was detected. These values are influenced by the choice of aggregation and clustering parameters, which can be adjusted to modulate sensitivity to smaller pocket-like regions. Here, we adopted the sc-PDB distance threshold to ensure consistency with established datasets. To assess PickPocket’s performance on predicted structures, we repeated the analysis using AlphaFold2-generated [37, 38] models of the same protein chains. Predictions were obtained for 74.8% of structures, with 98.9% overlap in 11 pocket detection between experimental and predicted conformations. This consistency suggests that PickPocket is robust to structural variations, making it applicable to both crystallographic and computationally derived protein models.

To assess the conservation and structural variability of binding pockets across evolutionarily related proteins, we analyzed three homologous superfamilies, selecting a representative structure for each family. We retrieved all available homologs for these reference proteins, predicted binding pockets using PickPocket, and superimposed the structures to evaluate pocket conservation based on centroid distances. For metalloproteases (TldD/PmbA, N-terminal domain), we selected 1VPB, a putative modulator of DNA gyrase (BT3649) from Bacteroides, and analyzed 11 homologous structures within the superfamily. For GPCRs (family 2, extracellular hormone receptor domain), we used 7UZO, the parathyroid hormone 1 receptor extracellular domain complexed with a peptide ligand, comparing it to 191 homologous structures. For kinases (protein kinase-like domain superfamily), we selected 6OQO, a CDK6 complex with an experimental inhibitor, and examined 8,000 homologs from the PDB. Binding pocket predictions were performed on the reference structures, while homologous proteins were superimposed to compute the distance between pocket centroids. Among aligned structures (defined as those with no missing atoms and at least 10 matching C*α* residues), the average RMSD between pocket centroids was 4.4 Å for metalloproteases, 5.92 Å for GPCRs, and 3.8 Å for kinases.

The large-scale assessment of PickPocket across experimental and predicted structures demonstrates its capacity to detect ligand-binding sites in diverse protein architectures. The high overlap between pocket predictions in crystallographic and AlphaFold2-derived structures suggests that PickPocket generalizes well across structural conformations, even in the absence of explicit holo-state training. This robustness is particularly relevant given the increasing reliance on computationally predicted structures in drug discovery, where experimental data may be unavailable. However, the small fraction of discrepancies highlights the potential influence of conformational flexibility, particularly in cases where ligand-induced pocket formation is essential. The analysis of homologous superfamilies further reveals how PickPocket captures both conserved and variable aspects of binding site topology. The lower RMSD observed for kinases compared to GPCRs aligns with known functional constraints, as kinases exhibit strong evolutionary pressure to maintain ATP-binding sites, whereas GPCR extracellular domains undergo more structural variation to accommodate different ligands. These findings suggest that PickPocket’s predictions are influenced by both structural rigidity and evolutionary conservation.

### 2.6 Predicted Pocket Embeddings Enable Binding Site Pairing

To assess whether pocket embedding similarity correlates with ligand compatibility, we docked 4,300 diverse compounds across selected receptor pairs and analyzed their embedding distances alongside ligand-binding distributions (See Appendixes C.2 and C.3). Our analysis revealed that pockets with lower embedding distances generally accommodate overlapping ligands, while those with higher distances show minimal shared ligand preferences.

Low-distance pocket pairs demonstrated high ligand-binding overlap. For example, 6FW0 _B – 2Z5Y _A (Monooxygenase-Monooxygenase, Distance = 6.04) and 8F07 _A – 8SKL _A (Hydrolase-Hydrolase, Distance = 14.82) showed strong ligand correlation. Interestingly, 7WCM _R – 6FW0 _A (GPCR-Monooxygenase, Distance = 13.73) exhibited unexpected ligand compatibility, suggesting conserved physicochemical properties. Most high-distance pairs showed minimal ligand overlap, though 8SKL _A – 7WCM _A (Hydrolase-G-Protein, Distance = 37.96) was a notable exception, showing strong ligand compatibility despite its high embedding distance. Further investigation of this case can be found in Appendix C.4.

These findings demonstrate that embedding similarity can guide target expansion, particularly for structurally related proteins. However, the results also indicate that additional descriptors may be needed to refine ligand-based predictions, especially when considering cross-family compatibility.

## 3 Conclusions

We present a method that recovers binding sites directly from sequence and structure features and that scales to proteome level analysis. The model integrates evolutionary signal with geometric context and yields consistent gains in residue level site recovery across diverse folds and ligand classes. In comparative evaluations it favours recall while maintaining competitive precision, which is advantageous when the aim is to identify plausible cavities for downstream design and screening.

Ablations show that evolutionary information carries substantial predictive power and that sequence representations complement but do not simply duplicate structural cues. This supports a view of binding as an evolved property that leaves a detectable imprint in multiple modalities. The method generalises across families and recovers residues that define volume, shape, and chemistry of pockets, which are the determinants that matter for docking and fragment growth rather than centroid proximity alone.

Large scale application produces maps that are useful beyond hit finding. Site calls are enriched in residues with functional annotation and with literature evidence of ligand engagement, which enables target assessment at the family level and rapid nomination of proteins with tractable surface chemistry. These maps can guide mutagenesis, help prioritise regions for cryo EM focused classification, and inform selection of constructs for biophysical assays.

There are limitations that set the agenda for next steps. Class imbalance and label noise in legacy datasets can bias calibration. Precision oriented variants sometimes produce higher Matthews correlation, and false positives tend to occur in shallow grooves and highly solvent exposed regions. Structural coverage still constrains the search space, and performance on proteins with extensive disorder remains weaker.

The broader implication is that site prediction should be evaluated on consequences for discovery decisions. When the goal is to avoid missing a true pocket, a recall leaning operating point is often the correct choice, especially when followed by inexpensive triage. When the goal is to annotate a small number of high confidence sites on a well studied target, a more conservative threshold may be preferable. Our framework supports both regimes and makes these trade offs explicit.

In summary, the method advances residue level binding site recovery in a way that is practical at scale and that connects to biological questions. It delivers interpretable maps that can shorten the path from structure to experiment and that can be integrated with variant data, dynamics, and ligand design. We believe that this approach can serve as a reliable front end for hit discovery and for systematic analysis of potential druggability across proteomes.

## 4 Methods

### 4.1 PickPocket

We propose PickPocket, a joint sequence-structure protein binding site prediction model that leverages serial fusion of ESM-2 and GearNet [39]. Serial fusion enables effective integration of evolutionarily conserved patterns from sequences with geometric structural relationships by using ESM-2’s contextual representations to initialize GearNet’s node features. This fusion strategy has been shown to outperform parallel and cross fusion approaches while maintaining architectural simplicity. By initializing GearNet with ESM-2’s pretrained sequence knowledge and using a reduced learning rate for ESM-2 to preserve its pretrained representations during training, our model effectively combines the complementary strengths of both sequence-based language modeling and structure-based geometric learning for accurate pocket identification.

#### 4.1.1 Sequence Representation with ESM-2

ESM-2 [23], a transformer-based protein language model, generates sequence embeddings for each residue in a protein sequence ℛ = [*r*_1_, *r*_2_, …, *r*_*n*_]. The sequence embedding process begins by encoding each residue into an initial feature vector:

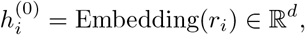

where *d* is the embedding dimension. Through a stack of transformer layers, these embeddings are refined using multi-head self-attention:

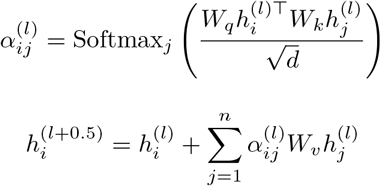

The final layer produces sequence embeddings *h*^(*L*)^ that capture contextual and evolutionary information.

#### 4.1.2 Structure Representation with GearNet

GearNet [24] processes the protein structure as a graph *G* = (*V*, ℰ, ℛ), where nodes represent residues and edges define their relationships. We establish three types of edges: sequential edges connecting residues within distance of 3 in the primary sequence, spatial edges connecting residues within 10Å, and k-nearest neighbor edges (k=10). Following serial fusion, each node is initialized with its corresponding ESM-2 sequence embedding:

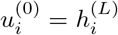

The structural information is integrated through 6 layers of message passing networks:

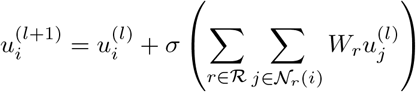

where *N_r_*(*i*) represents neighbors of node *i* connected by edge type *r*. The network incorporates batch normalization and residual connections throughout these layers.

### 4.2 Binding Site Prediction Architecture

#### 4.2.1 Model Initialization

ESM-2 is initialized with its pretrained weights from large-scale protein sequence data, while GearNet is initialized with the weights from residue type prediction [39]. This pretraining approach allows both models to leverage their respective pretrained knowledge of sequence and structure.

#### 4.2.2 Residue-Level Classification

The final residue representations are created by concatenating outputs from both ESM-2 and GearNet. These are fed into a two-layer multilayer-perceptron (MLP) [40] classifier:

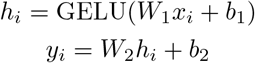

where *y*_*i*_ represents the predicted binding probability for residue *i*.

#### 4.2.3 Pocket Extraction

After obtaining residue-level predictions, we employ DBSCAN [41] clustering to identify cohesive binding pockets. Residues with binding probabilities exceeding 0.5 are considered for clustering, using parameters eps = 5Å (maximum distance between two residues to be considered neighbors) and min samples = 3 (minimum number of residues required to form a cluster). This post-processing step helps identify contiguous regions that are likely to form functional binding sites.

### 4.3 Training-testing Strategy

#### 4.3.1 Training Process

The training process jointly optimizes all components of the model using different learning rates: a lower learning rate of 10^−5^ for the ESM-2 parameters, and 10^−4^ for both GearNet and MLP classifier parameters. The model is trained using a smooth F1 loss function:

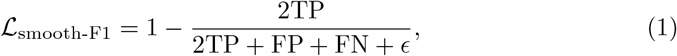

where TP, FN, and FP represent true positives, false negatives, and false positives respectively. We use the Adam optimizer with a batch size of 4. The smooth F1 loss helps balance the treatment of positive and negative cases, making it particularly suitable for imbalanced datasets.

While center-based distance metrics such as Top-N (also referred to as Distance from Center to Center (DCC)) are common, they are misleading for binding-site evaluation as they collapse pockets to a single point, reward arbitrary centroid proximity, and are highly sensitive to small translations that do not affect which cavity is actionable. Because screening and docking depend on which residues define the cavity, its extent, shape, and chemistry, rather than the precise centroid, we adopt a residue-level objective and optimize a differentiable F1 loss.

### 4.4 Datasets

#### Training

The sc-PDB (Structural Chemogenomics Protein Data Bank) [13, 42] is a curated dataset of druggable protein-ligand complexes extracted from the PDB. Compared to datasets like PDBBind [43] and Binding MOAD [44, 45], sc-PDB emphasizes druggable sites and ligand diversity, making it a valuable resource for computational drug design. The dataset used for training and validation is the 2017 release of sc-PDB database [13], which comprises 17,594 structures, 16,034 entries, 4,782 proteins, and 6,326 ligands. We used a subset of this dataset following PUResNet [11] and EquiPocket [46], where structures were clustered based on their Uniprot [26] IDs, and protein structures with the longest sequences were selected from each cluster. The training split follows VN-EGNN but with two additional constraints: proteins longer than 1,022 residues were excluded due to ESM embedding limitations, and the dataset was restricted to single-chain proteins only. These preprocessing steps resulted in a final dataset of 3,520 protein chains for training and validation.

#### Testing

We utilized the LIGYSIS dataset [25], a curated resource for evaluating protein–ligand binding site prediction methods. LIGYSIS comprises approximately 30,000 biologically relevant protein–ligand complexes, aggregating unique ligand-binding interfaces across multiple structures of the same protein. The dataset includes diverse ligand types—ions, peptides, nucleic acids, and small molecules—accounting for approximately 40% ion-binding sites, which are largely absent in other benchmarks. LIGYSIS aggregates data from multiple structures of the same protein, offering a comprehensive view of ligand-binding diversity.

For instance, human pancreatic alpha-amylase, represented by PDB entry 4GQQ in PDBbind, is expanded in LIGYSIS to include 13 unique binding sites derived from 51 structures. This approach significantly enhances dataset diversity, as indicated by a Shannon entropy of 8.8 in the ion-excluded subset (LIGYSIS_NI_), surpassing all other benchmark datasets. Compared to sc-PDB_FULL_, which limits each protein to the most relevant ligand, LIGYSIS provides a richer representation of binding site diversity.

Unlike prior datasets such as sc-PDB_FULL_, bMOAD_SUB_, CHEN11, PDBbind_REF_, SC6K, HOLO4K, and COACH420, LIGYSIS considers biological units instead of crystallographic asymmetric units, ensuring that only functional macromolecular assemblies are represented. Redundant protein–ligand interfaces, often inflated because of symmetry in crystallographic data, were systematically removed by clustering ligand interaction sites based on their protein interaction fingerprints.

LIGYSIS diverges from HOLO4K and COACH420, which rely on asymmetric units, by avoiding redundancy through a focus on biologically meaningful assemblies, improving the reliability of benchmarking results. Additionally, it corrects overrepresentation in datasets such as SC6K by ensuring non-redundant multimeric interactions and ligand diversity.

The final LIGYSIS benchmark set, after removing chains with missing residue mappings to UniProt, comprises 2,775 protein chains. To accommodate the maximum input size for ESM-2 embeddings, we crop proteins longer than 1022 residues to the first 1022 residues. Residues that were cropped are excluded from the ground truth. This preprocessing step led to the exclusion of 7 pockets in total from the LIGYSIS dataset.

## Appendix A Ablation Studies

**Table A1:** Performance Metrics for Different Ablation.

| Ablation | F1 Score | MCC | Top-N (%) | Top-N+2 (%) |
| --- | --- | --- | --- | --- |
| Freeze ESM | 0.35 | 0.32 | 37.5 | 40.3 |
| Freeze GearNet | 0.40 | 0.35 | 47.3 | 51.5 |
| No Freezing | 0.42 | 0.37 | 48.9 | 53.3 |

To assess the relative contributions of sequence-based and structure-based features in PickPocket, we conducted an ablation study by systematically freezing different components of our model during training. Specifically, we evaluated three conditions: (i) freezing the ESM embeddings while allowing GearNet and the classification head to update, (ii) freezing the GearNet encoder while updating ESM embeddings and the classifier, and (iii) training all components without freezing any parameters. The results of this experiment are summarized in Table A1.

When freezing ESM, the model achieved an F1 score of 0.35, an MCC of 0.32, and a Top-N recall of 37.5%, the lowest among all tested configurations. This decline in performance suggests that evolutionary embeddings from ESM provide essential residue-level contextual information that significantly enhances binding site prediction. Without updating these embeddings, the model primarily relies on structural features extracted by GearNet, which alone are insufficient for maximizing prediction accuracy. The particularly low Top-N and Top-N+2 recall further indicate that the model struggles to consistently rank correct binding sites among the top candidates when sequence-derived information is not actively incorporated.

Freezing GearNet while allowing ESM embeddings to update resulted in improved performance compared to freezing ESM. The F1 score increased to 0.40, and the MCC reached 0.35, while Top-N recall improved to 47.3%. This result suggests that while structural representation learning is beneficial, evolutionary embeddings alone capture a substantial portion of the predictive signal. The relatively minor performance drop compared to the fully trainable model highlights the robustness of ESM’s pretrained features, which retain essential sequence-derived patterns even when the structural encoder remains static. However, the lower Top-N recall compared to the no-freezing condition implies that structural refinement via GearNet enhances the model’s ability to prioritize true binding pockets.

Allowing both ESM and GearNet to update during training yielded the highest overall performance, with an F1 score of 0.42, an MCC of 0.37, and a Top-N recall of 48.9%. This configuration demonstrated the strongest ability to correctly rank binding sites and identify residue-level features indicative of ligand interaction. The improvements in Top-N and Top-N+2 recall indicate that integrating both evolutionary and structural information leads to more reliable predictions. These results confirm that PickPocket benefits from joint optimization of sequence and structure representations, where evolutionary embeddings guide feature extraction while geometric learning refines residue interactions.

The ablation study highlights the complementary nature of protein language models and geometric graph learning for binding site prediction. ESM embeddings provide deep contextual insights derived from large-scale evolutionary training, enabling the model to recognize conserved functional motifs that may not be immediately apparent from structural data alone. On the other hand, GearNet captures local residue-residue interactions, geometric constraints, and physicochemical properties, refining predictions based on structural context.

Our findings suggest that sequence features alone provide a strong baseline for binding site identification, as demonstrated by the relatively high performance of the freezing GearNet embeddings ablation. However, the inclusion of structural learning improves recall and pocket ranking, demonstrating the added value of geometric deep learning in fine-tuning binding predictions. Conversely, removing trainable sequence embeddings weakens the model’s ability to generalize, reinforcing the necessity of evolutionary signals in identifying functionally relevant sites.

## Appendix B Differential Embedding Similarity of Cryptic Pocket Residues

To investigate how sensitive PickPocket is to conformational changes, we analyzed the similarity of residue-level embeddings between apo and holo states using *centered kernel alignment (CKA)*. Four apo holo protein pairs were considered: 1AKE _A - 4AKE _A, 2PC0 _A - 4EJL _A, 2W9T _A - 2W9S _A, and 4P0I _B - 5OTA _B. For each pair, embeddings were extracted separately for pocket and non-pocket residues. Pocket and non-pocket aminoacids were first matched between apo and holo structures. Embeddings were aligned based on these common residues, and CKA was computed between apo and holo embeddings using the centered formulation:

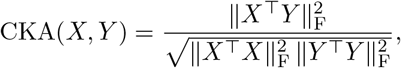

where *X* and *Y* are mean-centered embedding matrices and ∥ · ∥_F_ denotes the Frobenius norm.

The analysis was performed separately for pocket (CKA_site_) and non-pocket (CKA_non_) residues, and their difference ΔCKA = CKA_site_ – CKA_non_ was computed for each protein pair. The obtained values were:

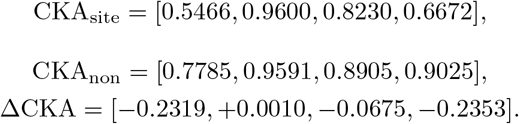

Across the four protein pairs, CKA_site_ values were generally lower than CKA_non_, yielding a mean ΔCKA of **–0.133 ± 0.103**. Non-pocket residues thus exhibited higher representational similarity between apo and holo states, reflecting their structural stability, while pocket residues showed larger embedding shifts consistent with conformational or functional rearrangements upon ligand binding. These findings highlight that PickPocket encodes dynamic and functionally relevant variations in pocket regions, demonstrating sensitivity to small but biologically important conformational changes.

## Appendix C Pairing binding sites with PickPocket embeddings

### C.1 Docking Protocol

We used the Glide [47] software to perform SP docking on the selected structures. In most cases, we positioned the grid by selecting the nearest point to the center of mass based on predictions from PickPocket. In certain instances, we included additional residues to define a more meaningful docking region for the compounds.

### C.2 Diversity dataset

We selected compounds from the ZINC22 database [48] based on specific physicochemical properties and availability. The filtering criteria were:

- Molecular weight (MW) *<* 425 Da
- LogP < 5
- In stock as of 2022 (when the dataset was downloaded)

This initial filtering yielded approximately 9 million compounds. To obtain a representative subset, we performed stratified sampling based on molecular weight. From this sampled set, additional manual selection was conducted to ensure structural diversity.

### C.3 Selected pairs and euclidean distances

To investigate the relationship between pocket embedding similarity and functional binding site overlap, we selected a diverse set of protein chain pairs and computed their Euclidean distances in the PickPocket embedding space. This analysis aimed to assess whether lower embedding distances correlate with greater ligand-binding site similarity, potentially enabling target expansion and ligand repurposing strategies.

We selected protein chains from distinct structural and functional categories to include a variety of ligand-binding domains. Pairs were chosen based on:

- Protein functional annotation from UniProt and PDB metadata.
- Presence of at least one PickPocket-predicted binding site with a confidence score above 0.5.
- Structural classification based on SCOP/CATH domains to ensure representation across different protein families.
- Diversity in molecular function, spanning hydrolases, monooxygenases, G-protein coupled receptors (GPCRs), and non-binding stabilizers (NB-stabilizers).

For each selected protein pair, we extracted the PickPocket-generated pocket embeddings and computed their Euclidean distances to quantify binding site similarity. The process involved the following steps:

1. **Pocket Representation:** For each protein chain, we identified the top-ranked predicted pocket based on PickPocket’s binding probability scores.
2. **Feature Extraction:** The pockets were represented as 4352-dimensional embeddings derived from the final layer of the PickPocket model, averaging the representation.
3. **Distance Calculation:** The Euclidean distance between the pocket embeddings of two protein chains was computed as:

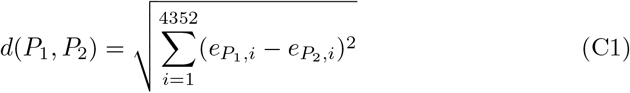

where 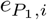 and 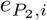 represent the *i*-th feature in the respective pocket embeddings.
4. **Functional Comparison:** The functional annotations of the protein chains were retrieved from UniProt, and their biological roles were compared to examine potential ligand-binding overlap.

Table C2 presents the computed Euclidean distances for selected protein pairs alongside their functional annotations. The distance values range from 6.04 (highly similar monooxygenases) to 37.96 (divergent hydrolase-G-Protein pair). Low embedding distances generally corresponded to functionally related proteins, supporting the hypothesis that PickPocket embeddings capture meaningful binding site relationships. However, certain high-distance pairs (e.g., 7WCM A – 8SKL A) exhibited unexpected ligand compatibility.

**Table C2:** Chain Pairwise Scores and Functional Annotations.

| Chain 1 | Chain 2 | Distance | Function 1 | Function 2 |
| --- | --- | --- | --- | --- |
| 6fw0_B | 2z5y_A | 6.04 | monooxygenase | monooxygenase |
| 8f07_A | 8skl_A | 14.82 | hydrolase | hydrolase |
| 6fw0_B | 7wcm_N | 37.07 | monooxygenase | nb-stabilizer |
| 8skl_A | 7wcm_A | 37.96 | hydrolase | G-protein |
| 7wcm_R | 6fw0_A | 13.73 | gpcr | monooxygenase |
| 6fw0_B | 8f07_A | 20.28 | monooxygenase | hydrolase |
| 6fw0_B | 8skl_A | 28.06 | monooxygenase | hydrolase |
| 2z5y_A | 7wcm_A | 31.78 | monooxygenase | G-protein |
| 6fw0_A | 8skl_A | 19.21 | monooxygenase | hydrolase |

### C.4 Results for 8SKL_A – 7WCM A

After superimposing the structures, we found that the high distance was influenced by a poor selection of residues from PickPocket. While these residues were near the binding site, they were insufficient to provide the resolution needed for accurate representations. Additionally, in the docking distance plot, we observe that the compounds contributing the most to the low distance in scores are those with high (closer to zero) docking scores. This suggests that even though the pocket exists in these pairs, the ligands tend not to dock with enough meaningful contacts to obtain a better score.

**Fig. C1:**
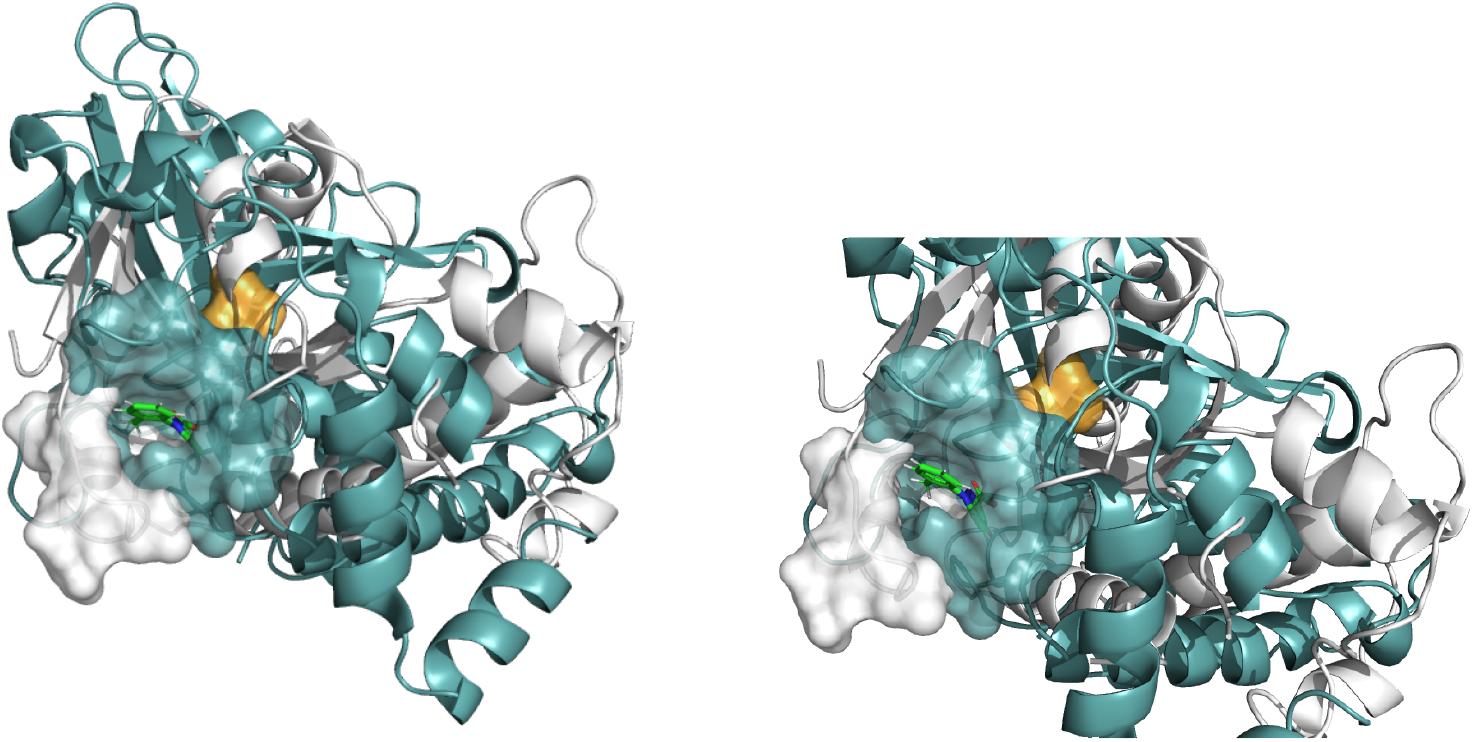
Pocket superposition of structures 8SKL_A (blue) and 7WCM_A (white). White surface indicates the pocket obtained from PickPocket prediction and in orange the one added to successfully perform docking.

**Fig. C2:**
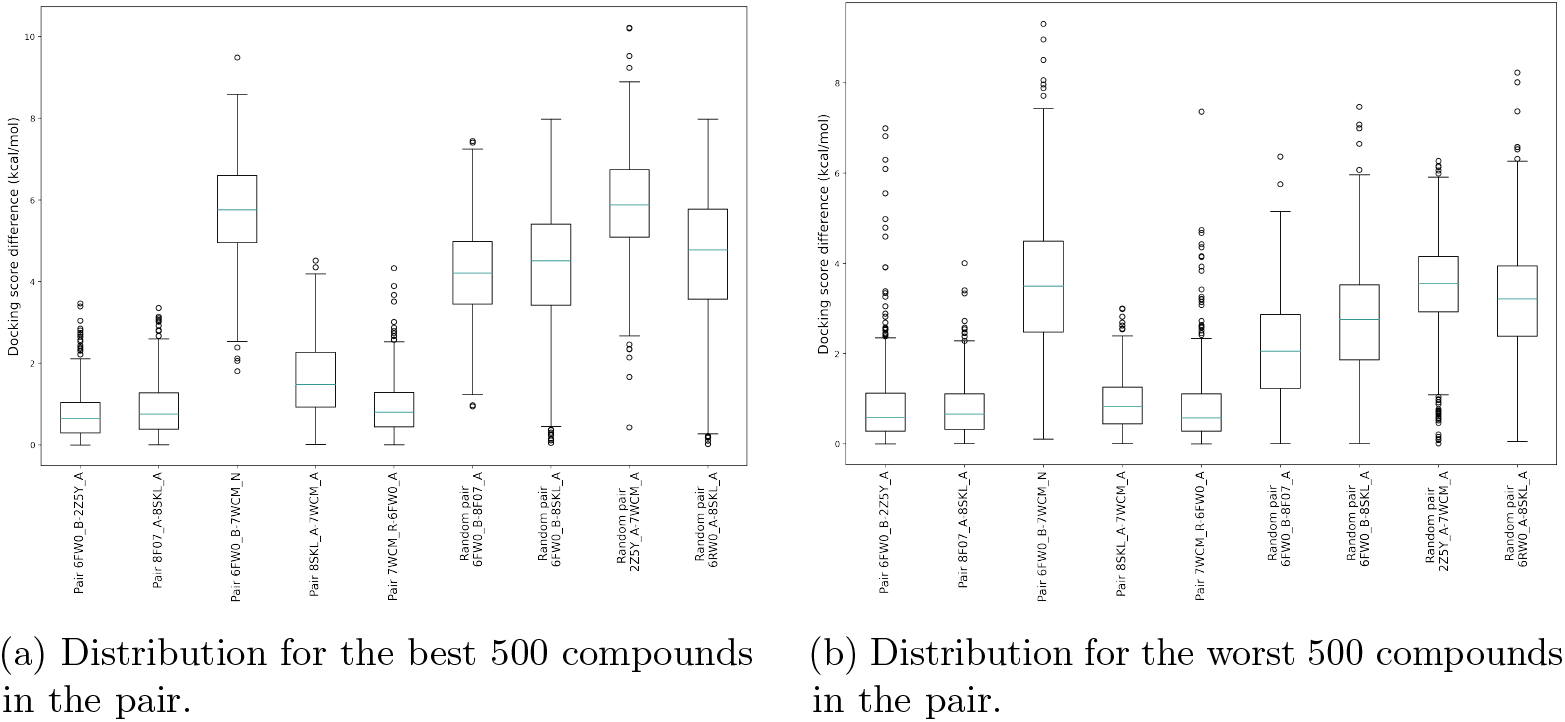
Distribution of docking score differences for the same compound across selected pairs. We evaluate this using the top 500 and bottom 500 compounds based on docking scores.

## Footnotes

1 Serial and Parallel implementations differ from the main implementation used in this work such as GearNet model is separately pretrained on Alphafold without being given ESM features. PickPocket leverages GearNet pretraining over AlphaFold database with knowledge of ESM embeddings.

